# SARS-CoV-2 activates ER stress and Unfolded protein response

**DOI:** 10.1101/2021.06.21.449284

**Authors:** Livia Rosa-Fernandes, Lucas C. Lazari, Janaina Macedo da Silva, Vinicius de Morais Gomes, Rafael Rahal Guaragna Machado, Ancely Ferreira dos Santos, Danielle Bastos Araujo, João Vitor Paccini Coutinho, Gabriel Santos Arini, Claudia B. Angeli, Edmarcia E. de Souza, Carsten Wrenger, Claudio R. F. Marinho, Danielle B. L. Oliveira, Edison L. Durigon, Leticia Labriola, Giuseppe Palmisano

## Abstract

Coronavirus disease-2019 (COVID-19) pandemic caused by the SARS-CoV-2 coronavirus infection is a major global public health concern affecting millions of people worldwide. The scientific community has joint efforts to provide effective and rapid solutions to this disease. Knowing the molecular, transmission and clinical features of this disease is of paramount importance to develop effective therapeutic and diagnostic tools. Here, we provide evidence that SARS-CoV-2 hijacks the glycosylation biosynthetic, ER-stress and UPR machineries for viral replication using a time-resolved (0-48 hours post infection, hpi) total, membrane as well as glycoproteome mapping and orthogonal validation. We found that SARS-CoV-2 induces ER stress and UPR is observed in Vero and Calu-3 cell lines with activation of the PERK-eIF2α-ATF4-CHOP signaling pathway. ER-associated protein upregulation was detected in lung biopsies of COVID-19 patients and associated with survival. At later time points, cell death mechanisms are triggered. The data show that ER stress and UPR pathways are required for SARS-CoV-2 infection, therefore representing a potential target to develop/implement anti-CoVID-19 drugs.

## INTRODUCTION

Coronavirus Disease 19 (COVID-19) is caused by severe acute respiratory syndrome coronavirus 2 (SARS-CoV-2) ^1^, an enveloped RNA virus belonging to the family *Coronaviridae* in the subfamily *Orthocoronavirinae* ^2^. Common symptoms of human infection are dry cough, sore throat and fever; however, a percentage of the patients can develop organ failure, septic shock, pulmonary edemas, severe pneumonia and Acute Respiratory Distress Syndrome, complications that can be fatal ^3^. Considering the fast increase in the infection numbers and the outbreaks of SARS-CoV-2 in other countries, on 30^th^ January of 2020 the World Health Organization (WHO) declared COVID-19 to be a Public Health Emergency of International Concern and warned that countries with vulnerable health care systems would be at high risk ^3^.

Understanding host-pathogen interactions and the host response to viral infection are important to develop new strategies to treat, prevent and diagnose COVID-19 ^4^. The host-pathogen dynamics is the key to infection control and minimize spread, incidence, prevalence and mortality ^5–8^. In the host cell, viral proteins are processed through the endoplasmic reticulum (ER) and Golgi apparatus shaping the glycosylation level (especially N-linked glycans) of each site and regulating their folding^9^. This post-translational modification is often used by viruses to evade immune recognition, to increase receptor binding, infectivity, viral release, virulence and to increase viral replication ^10–12^. Therefore, glycosylation process is the subject of numerous studies and often used as therapeutic target to treat viral infections ^13^. One strategy is to target the host glycosylation machinery to pharmacologically disrupt viral glycoproteins folding, being the inhibitors of N-linked glycosylation one of the most tested agents for antiviral use ^14^. In particular, recent reports have shown that not only the targeting of host machinery but a direct modification in glycosylation levels of viral glycoproteins could impair viral infection/replication of SARS-Cov-2, thus indicating that targeting this process is a promising strategy to reduce SARS-CoV-2 infection.

Viral infections typically lead to an increase in protein synthesis that can overwhelm the ER folding capacity, which may result in unfolded protein accumulation resulting in ER stress ^15^. To reduce this type of stress, the cell activates signaling pathways known as unfolded protein response (UPR), that reduces the overall protein synthesis, increases ER’s folding capacity and targets misfolded proteins to proteasome degradation ^16^. UPR consists of three signaling pathways activated by the transmembrane protein sensors IRE1, PERK and ATF6.

Briefly, IRE1 branch activation causes the mRNA splicing of a potent transcription factor, XBP1, which induces the expression of genes that will act in ER stress response. The first step in the PERK branch activation includes the increase on its phosphorylation state, which promotes the release of ER chaperone BiP as well as the phosphorylation of the transcriptional factor eIF2α, which will then upregulate ATF4 expression. This signaling pathway culminates with protein synthesis attenuation and selective induction of translation of ER chaperones and UPR-related transcriptional factors. Finally, the accumulation of unfolded proteins causes ATF6 release from the ER membrane allowing its traffic to the Golgi apparatus, where it will be activated by cleavage and consequently lead to upregulation of genes encoding for ER chaperones and components necessary for degradation of unfolded proteins ^17^.

Recognizing the biomolecular features that facilitate infection and which host-mediated mechanisms the pathogen uses to favor its replication and transmission are substantial to achieve disease control and prevention^18^. Quantifying and analyzing the temporal changes in host and viral proteins over the biological processes of infection could provide valuable information about the virus-host interplay ^19^. Here we applied a temporal and spatial proteome analysis combined with assessment of N-deglycoproteome to depict the host response to SARS-CoV-2 infection. We demonstrated that SARS-CoV-2 induces ER stress response, UPR and modulation of glycosylation machinery in the host cell. ER-associated transcripts upregulation was also detected in lung biopsies of COVID-19 patients and associated with higher survival. We also show that sustained infection prolonged the effects of ER-stress and UPR, leading to cell death related to necroptosis and caspase induced apoptosis pathways.

## MATERIALS AND METHODS

### Cell lines, SARS-CoV-2 and infection assays

Vero cell line (ATCC CCL-81) were maintained in DMEM medium supplemented with 10% (v/v) FBS, 4.5 g/L glucose, 2 mM L-glutamine, 1 mM sodium pyruvate, 100 U/mL penicillin-streptomycin and 1.5 g/L NaHCO3 at 37°C with 5% CO2. Calu-3 cells (ATCC HTB-55) were maintained in DMEM medium supplemented with 20% (v/v) FBS, 1% (v/v) nonessential amino acids, 4.5 g/L glucose, 2 mM L-glutamine, 1 mM sodium pyruvate, 100 U/mL penicillin-streptomycin and 1.5 g/L NaHCO3 at 37 °C with 5% CO2.

SARS-CoV-2 isolate (HIAE-02: SARS-CoV-2/SP02/human/2020/BRA (GenBank accession number MT126808) ^20^ was used to infect Vero CCL-81 and Calu-3 cells with multiplicity of infection (MOI) of 0.02. Following adsorption in DMEM with 2.5% FBS for 1h, fresh medium was added, and cells were further incubated at 37 °C and 5% CO2 for different time points (2, 6, 12, 24 and 48h). After the designated incubation time, cell lysates were retrieved in 1% sodium deoxycholate (SDC) in phosphate buffered saline solution with Protease Inhibitor Cocktail (cOmplete, Roche) buffer, 0.1M Na2CO3 with Protease Inhibitor Cocktail (cOmplete, Roche) buffer or 8M Urea with Protease Inhibitor Cocktail (cOmplete, Roche) buffer, according to the follow-up application. Aliquots of cells and supernatants were collected at the different time points for virus RNA copy number quantification by reverse transcription-quantitative polymerase chain reaction (RT-qPCR), targeting the E gene ^21^. The assay was reproduced in two independent experiments and expressed by standard error of the mean (SEM). Graphics and SEM were done using GraphPad Prism software version 8.1 (GraphPad Software, San Diego, USA).

All assays were conducted in triplicates in a BSL-3 facility at the Institute of Biomedical Sciences, University of Sao Paulo, under the Laboratory biosafety guidance related to coronavirus disease (COVID-19): Interim guidance, 28 January 2021 (https://www.who.int/publications/i/item/WHO-WPE-GIH-2021.1).

### Total proteome analysis (cell lysis and trypsin digestion)

SARS-CoV-2-infected and mock-infected control cells were lysed in 1% sodium deoxycholate (SDC), 1× PBS and 1× protease inhibitor cocktail (Sigma-Aldrich) and probe tip sonicated for three cycles for 20 s and intervals of 30 s on ice ^22^. Proteins were reduced with 10 mM DTT for 30 min at 56 °C and alkylated with 40 mM IAA for 40 min at room temperature, in the dark. Proteins were quantified using NanoDrop 2000 spectrophotometer (Thermo Scientific) before sequencing grade porcine trypsin (Promega) was added to a 1:50 ratio. The digestion, which proceeded for 16 h at 37 °C, was blocked by adding TFA 1% (v/v) final concentration before the SDC was removed from the solution by centrifugation at 10000 x g for 10 min ^22^. Tryptic peptides were desalted using reversed phase C18 microcolumns before LC-MS/MS analysis.

### Microsomal membrane proteome analysis (cell lysis and trypsin digestion)

Microsomal membrane protein fraction was isolated as previously described ^23,24^. Briefly, cells were lysed in 100 mM Na_2_CO_3_, pH 11 containing protease inhibitors cocktail (Sigma-Aldrich) by sonication using three rounds of probe-tip sonication at 40% output for 20 s with 30 s resting on ice. The lysates were incubated at 4°C with gentle rotation for 1.5 h followed by ultracentrifugation at 100000 x g for 1.5 h. After ultracentrifugation, the pellets were recovered and washed with 100mM triethylammonium bicarbonate (TEAB) and re-dissolved in 8 M urea in 50mM TEAB. Protein concentration determination was performed using Qubit fluorescent assay (Invitrogen). The solubilized membrane pellets were reduced and alkylated as described above. Urea was diluted to 0.8 M with 50mM TEAB and proteins were digested with trypsin at an enzyme to substrate ratio of 1:50 for 16 h at room temperature. Tryptic peptides were purified using Oligo R3 reversed phase SPE micro-column ^25^.

### Glycopeptide enrichment and PNGase F deglycosylation

Tryptic glycopeptides obtained from microsomal membrane proteins were enriched using HILIC SPE as previously described ^26,27^. Briefly, dried peptides were reconstituted in 100 μL loading and washing buffer containing ACN 80% (v/v) in TFA 1% (v/v). Peptides were loaded onto a primed custom-made HILIC SPE micro-column packed with PolyHYDROXYETHYL A™ resin (PolyLC Inc). The HILIC SPE columns were then washed in 100 μL loading and washing buffer. The enriched glycopeptides were eluted with TFA 1% (v/v) followed by 25 mM NH4HCO3 and finally ACN 50% (v/v). The three eluted fractions were then combined, dried by vacuum centrifugation and purified on a primed Oligo R3 reversed phase SPE micro-column. The enriched glycopeptides were resuspended in 50 mM Ambic, pH 7.5 and de-N-glycosylated using 500 U N-glycosidase F (PNGase F, New England Biolabs) for 12 h at 37°C. After incubation, the de-N-glycosylated were purified on a primed Oligo R3 reversed phase SPE micro-column, before LC-MS/MS analysis.

### LC-MS/MS proteomics analysis

Tryptic peptides were analyzed by nanoflow LC-MS/MS analysis. The nLC-MS/MS analysis was performed using an Easy nano LC1000 (Thermo) HPLC coupled with an LTQ Orbitrap Velos (Thermo). Peptides were loaded on a C18 EASY-column (2cm x 5 μm x 100 μm; 120 Å pore, Thermo) using a 300 nL/min flow rate of mobile phase A (0.1% formic acid) and separated in a C18 PicoFrit PepMap (10 cm x 10 μm x 75 μm; 135 Å pore, New Objective), over 105 minutes using a linear gradient 2-30 % followed by 20 min of 30-45% of mobile phase B (100% ACN; 0,1% formic acid). The eluted peptides were ionized using electrospray. The top 20 most intense precursor ions with charge-state ≥ 2 were fragmented using CID at 35 normalized collision energy and 10 ms activation time. The MS scan range was set between 350-1800 m/z, the MS scan resolution was 60.000, the MS1 ion count was 1×10e6 and the MS2 ion count was 5×10e4. The mass spectrometry proteomics data have been deposited to the ProteomeXchange Consortium (http://proteomecentral.proteomexchange.org) via the PRIDE partner repository ^28^.

### Database Search and Statistical Analysis

Raw data were searched using Proteome Discoverer computational platform v2.3.0.523 (PD) using the Sequest search engine. The parameters used for database search were *Chlorocebus* (20,699 entries downloaded on 12/2020) proteome databases supplemented with the UniProt SARS-CoV-2 proteome and with the common contaminants. Trypsin as cleavage enzyme, two missed cleavages allowed, carbamidomethylation of cysteine as fixed modification, oxidation of methionine, and protein N-terminal acetylation as variable modifications. Asparagine and glutamine deamidation were included as variable modifications in the de-glycoproteome data. In the Proteome Discoverer platform, the percolator, peptide, and protein validator nodes were used to calculate PSMs, peptides, and proteins FDR, respectively. FDR less than 1% was accepted at protein level. Protein grouping was performed using the strict parsimony principle. Label-free quantification was performed using the extracted ion chromatogram area of the precursor ions. Protein quantification normalization and roll-up were performed using unique and razor peptides and excluding modified peptides. Differentially regulated proteins between the three conditions were selected using *t test* with a post-hoc background-based adjusted p-value <0.05 for multiple hypothesis correction ^29^.

### Bioinformatics analysis

Gene ontology (GO) was performed using the g: profiler tool and GOplot package ^30^, available in Bioconductor. A q-value threshold of 0.05 was used, corrected by the Benjamini-Hochberg method ^31^. Enriched pathways were determined by Reactome and KEGG platform (q-value < 0.05)^32^; complementary analyses were performed using the ReactomeFIPlugIn app ^31^. Protein’s subcellular locations were determined by UniProt release 12.4 (https://www.uniprot.org/news/2007/10/23/release). The “Peptides” package ^33^ was used to determine the hydropathy score of glycopeptides and the “mixOmics” package ^34^ was used to integrate the data for total, membrane, and deglycoproteome. Complementary analyses were performed using Perseus, ggplot2 package, Graphpad prism v.8, and RStudio software.

### Structural analysis of identified peptides

Structural data for full-length SPIKE protein was retrieved from the CHARMM-GUI coronavirus repository, based on the model of Wrapp et al ^35^, while ORF8 protein structure was downloaded from PDB ^36^. Peptides identified through MS data analysis were searched in the protein to better visualize their regions using PyMOL 2.4.1

### Western blotting

Cells were lysed in SDC buffer containing protease (Roche, Basel, Switzerland) and phosphatase (Sigma-Aldrich) inhibitor cocktails. Proteins (10ug of each cell lysate) were separated by SDS-PAGE and electro transferred onto PVDF membranes. They were subsequently blocked in a solution containing 5% milk in PBS-Tween 0.1% (v/v) for 1h at room temperature (RT). Primary antibodies were diluted in the blocking solution (**Table 1**) and were incubated overnight at 4°C. Membranes were washed three times in PBS-Tween (0.1%) and then incubated at RT for 1h with HRP-labeled secondary antibodies, diluted in a solution of 0,1% BSA in PBS-Tween 0.1% (v/v). Monoclonal anti-alpha-tubulin clone B-5-1-2 antibody (T5168, Sigma-Aldrich) was used as the loading control. For phospho-protein quantification, the membranes were stripped, blocked and reprobed using a solution containing the corresponding anti-fosfospecific antibody. Proteins were visualized by using enhanced chemiluminescence (Millipore Corporation, Billerica, MA, USA). Images were acquired using Uvitec Image System (Cleaver Scientific Limited, Cambridge, UK). Quantitative densitometry was carried out using the ImageJ software (National Institutes of Health). The volume density of the chemiluminescent bands was calculated as integrated optical density × mm2 using ImageJ Fiji. Phosphorylated proteins densitometry values were divided by the total protein values and the α-tubulin antibody was used as the normalizer of the amount of proteins applied in the gel. At least three independent experiments were performed for each cell type and condition.

**Table 1:** List of primary antibodies used for protein detection by Western blot.

| Protein | Company | Catalog | Dilution |
| --- | --- | --- | --- |
| Phospho(S345) MLKL | Abcam | ab196436 | 1:1000 |
| MLKL | Abcam | ab184718 | 1:1000 |
| GPX4 | Abcam | ab125066 | 1:1000 |
| NRF2 | Abcam | ab137550 | 1:1000 |
| Phospho-PERK (Thr980) | Cell Signaling | #3179S | 1:500 |
| PERK | Cell Signaling | #3192 | 1:1000 |
| phospho-eIF2α (S51) | Cell Signaling | #3597S | 1:1000 |
| eIF2α | Cell Signaling | #5324 | 1:1000 |
| ATF4 | Cell Signaling | #11815 | 1:1000 |
| phospho-IRE1α (S724) | Abcam | ab48187 | 1:1000 |
| IRE1α | Cell Signaling | #3294 | 1:1000 |
| XBP1s | Cell Signaling | #12782 | 1:1000 |
| ATF6 | Abcam | ab11909 | 1:1000 |
| CHOP | Cell Signaling | #2895 | 1:500 |
| BIP | Cell Signaling | #3183 | 1:1000 |
| Clivead-CASP3 (Asp175) | Cell Signaling | #9661L | 1:500 |
| CASP3 | Cell Signaling | #9668S | 1:500 |
| CASP9 | Cell Signaling | #9508S | 1:1000 |
| Alpha-tubulin clone B-5-1-2 | Sigma-Aldrich | T5168 | 1:10.000 |
| Anti-rabbit | Vector Laboratories | PI1000 | 1:1000 |
| Anti-mouse | Vector Laboratories | PI2000 | 1:1000 |

### Statistical analysis

All western blot results were analyzed for Gaussian distribution and passed the normality test (the number of independent experiments was chosen to present a normal distribution). The statistical differences between group means were tested by One-way ANOVA followed by Tukey’s post-test for multiple comparisons. A value of p<0.05 was considered as statistically significant in all analysis. Results are presented as mean ± S.E.M. Each dot represents an independent experiment.

### RNA-seq data reanalysis

The fastq files were downloaded from the https://sra-explorer.info/ platform with the BioProject accession number PRJNA646224 ^37^ and processed on the Galaxy server ^38^. The ‘FastQC’ module was used to report the quality reads, following by the trimmed using Trim Galore (v. 0.4.3.1) set to the single-end library. The Trim Galore output sequences were aligned to the human reference genome hg38 using the HISAT2 (Galaxy Version 2.1.0+galaxy7) platform. A count table was generated using the htseq-count (Galaxy Version 0.9.1). The differently regulated genes were analyzed by the limma, Glimma, edgeR, and Homo.sapiens packages applying a cut-off of |log2FC|>1 and a p-adjusted value <0.05 (Benjamini-Hochberg).

## RESULTS

To identify molecular pathways affected by viral-host interplay on the course of SARS-CoV-2 infection, a spatio-temporal MS-based quantitative approach comprised of proteome, membranome and N-deglycoproteome of SARS-CoV-2 infected Vero cells was conducted at 2, 6, 12 and 48 hpi. The membranome refers to the analysis of microsomal-enriched proteins while the N-deglycoproteome refers to the analysis of formerly N-linked glycopeptides associated proteins. Validation of differentially expressed proteins was performed in human epithelial lung cells (Calu-3) by western blotting and *in silico* transcriptome analysis of lung biopsies from COVID-19 patients and controls (**Figure 1A**).

**Figure 1.**
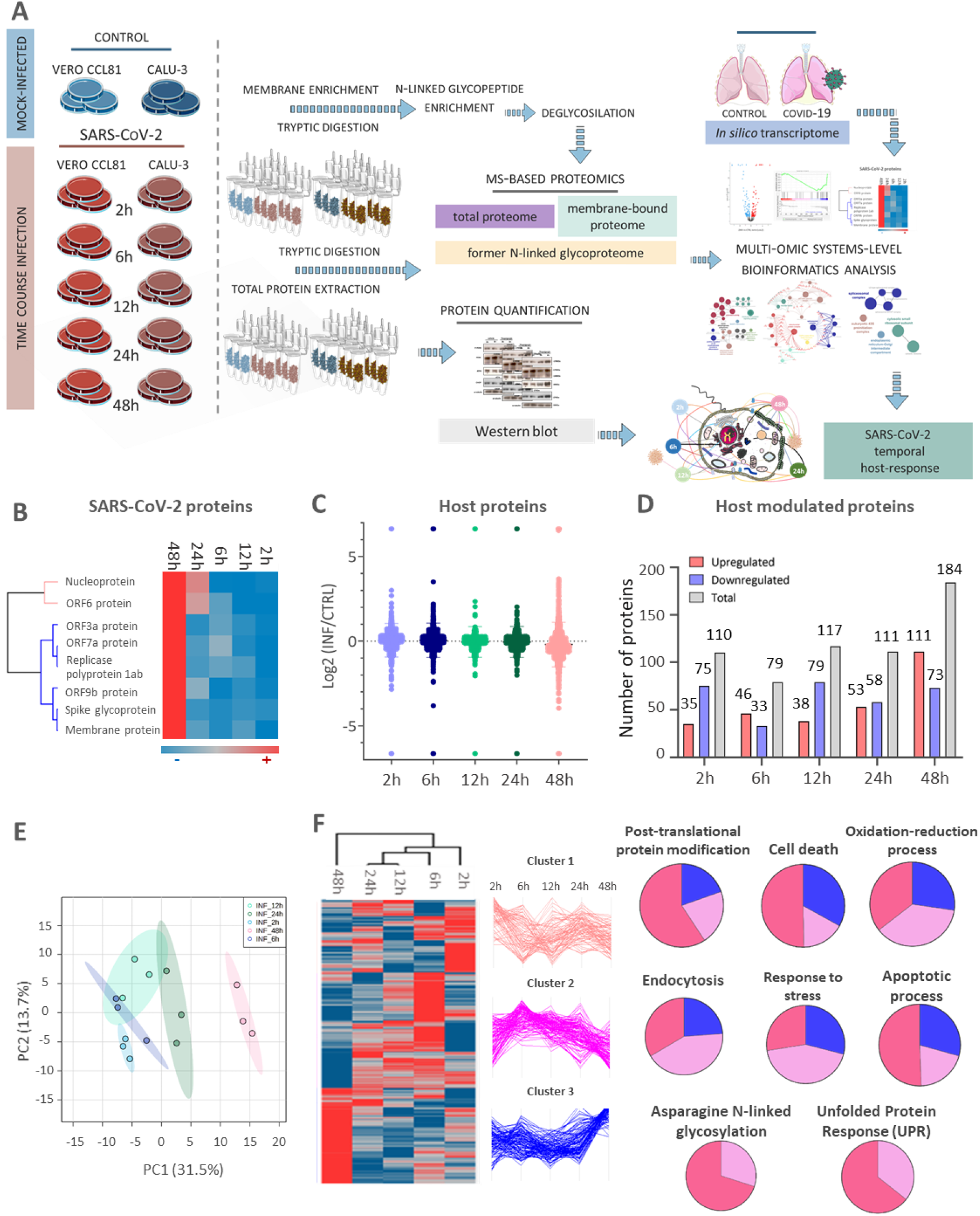
Time-resolved proteome modulation upon SARS-CoV-2 infection. Experimental approach applied to access spatiotemporal host-response to SARS-CoV-2 infection included evaluation of proteome, membranome and N-deglycoproteome of infected cells combined with WB protein quantification and transcriptome analysis of lung tissue of COVID-19 patients (A). Heatmap of SARS-CoV-2 viral proteins expression. Red and blue colors indicate high and low expression, respectively (B); Principal component analysis of quantitative proteome changes during infection(C); Quantitative proteome profile of Vero cells infected with SARS-CoV-2 after 2h, 6h, 12h, 24h, and 48h. The log2 ratio of infected vs control is shown (D); Vero cell proteins differentially regulated between the control (CTRL) and infected (INF) paired groups (q-value <0.05) at different time points. Red, blue and grey bars indicate up, down and total regulated proteins, respectively (E); Differentially regulated host proteins in at least one time point. Proteins were grouped into clusters associated with early, middle, and late events, respectively. Representation of enriched biological processes (BP) per cluster (q-value <0.05) (F).

A total of 1842 proteins were identified and quantified in the proteome analysis (**Supplementary Data S1**). Eight viral proteins were identified, being three structural proteins (M, S and N) and 5 non-structural proteins (ORF3a, ORF6, ORF9b, ORF7a and ORF1ab) (**Figure 1B, Supplementary Data S1**). These proteins increased over time showing a steeper upsurge already after 6 hpi, in agreement with the qPCR data (**Supplementary Figure 1**). Respectively, PCA analysis of quantitative host-proteome features showed a clear separation between early (2 and 6 hpi) and late (24 and 48 hpi) infection times (**Figure 1C**). Host proteome regulation varied across time, showing preponderant downregulation until 48 hpi, when most regulated proteins were up-regulated compared to control (**Figures 1D and E, Supplementary Data S1**). For the time-points of 2, 6, 12, 24 and 48 hpi, we identified a total of 110, 79, 117, 111 and 184 regulated proteins, respectively (**Figure 1E, Supplementary Data S1**).

To understand the global changes taking place in the host proteome during viral infection, we analyzed the differences in protein levels over time in a system-wide manner (**Figure 1F**). The host proteome already changed in the early time-points (2 and 6 hpi), but 48 hpi showed most extensive modulation. We identified three main clusters which contained proteins that participate in several key biological processes for early, middle and late times of host response expression profile (**Figure 1F**). Our analysis revealed processes related to post-translational protein modification, cell death, oxidation-reduction, endocytosis, response to stress, unfolded protein response (UPR), apoptosis and N-linked glycosylation (**Figure 1F, Supplementary Data S1**). Interestingly, assessment of protein subcellular location showed that while ER-related proteins more representative in the early time-points than at 48 hpi, the opposite pattern was observed for Nucleus-related proteins (**Figure 2A**). We found the expression of proteins associated with glycosylation biosynthesis modulated through the course of infection (**Figure 2B, Supplementary Data S1**). Of note are the ones related to nucleotide sugar biosynthesis, proteoglycans and glycosyltransferases (**Figure 2C)** besides intracellular membrane-bound organelle, endomembrane system and extracellular region cellular location **(Figure 2D).** Moreover, biological processes and pathways related to stress response and Asparagine N-linked Glycosylation were already observed at 2 hpi. In addition, alteration of proteins involved in UPR regulation were observed at 6 hpi (**Figure 2E and F**). These data indicate a remodeling of the host glycoproteome upon SARS-CoV-2 infection.

**Figure 2.**
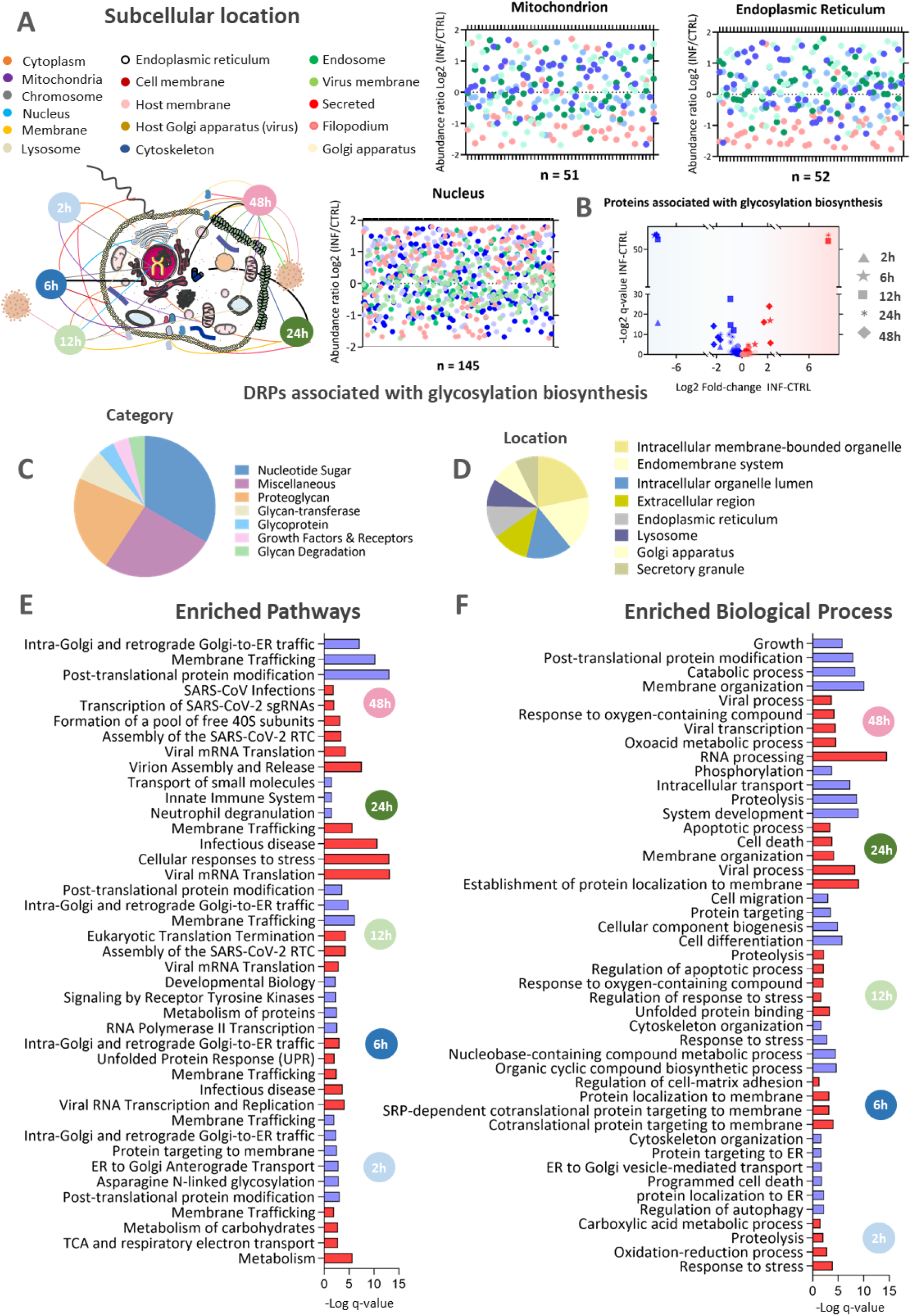
Time-resolved functional analysis of differentially expressed proteins upon SARS-CoV-2 infection. Abundance ratio (log2 infected vs control) of differentially regulated proteins associated to nucleus, mitochondria and ER according to infection time (A); Volcano plot of proteins associated to the glycosylation biosynthesis modulated in SARS-CoV-2-infected Vero cells vs control Up and downregulated proteins are represented in red and blue, respectively (B); Category (C) and subcellular location (D) of differently regulated proteins associated to the glycosylation biosynthesis. Enriched pathways (E) and biological processes (BP) (F) at 2, 6, 12, 24 and 48 hpi (q-value <0.05).

Since we observed glycosylation processes and multiple biological processes involving ER and membrane proteins, we proceeded with the evaluation of membranome and N-deglycoproteome (**Figure 3A, Supplementary Data Set 2**). We found 323 proteins identified in all three approaches, showing an increase in proteome coverage by enriching for glycosylated and membrane proteins (**Figure 3A**). As observed in the proteome, the number of regulated proteins in the membranome also increased over time, and at 48 hpi the number of down-regulated proteins was higher than the up-regulated ones (**Figure 3B**). Host proteins associated to glycoconjugate biosynthesis were mapped (**Figure 3C**). Moreover, we identified 1,037 N-deglycopeptides from the host (**Figure 3D, Supplementary Data Set 2**), being 545 regulated ones belonging to 338 N-deglycoproteins (**Supplementary Data Set 2**). The number of regulated N-deglycopeptides increased over time, but differently from the total proteome, most of which were downregulated at 48 hpi (**Figure 3E, Supplementary Data Set 2**). Expression pattern of the regulated N-deglycopeptides indicated a formation of three clusters (**Figure 3F, Supplementary Data Set 3**). In the first cluster an increase in the N-deglycopeptide abundance was observed over time while in cluster 3 there was a decrease. The hydropathy score associated to N-deglycopeptides in cluster 3 was significantly higher than the ones in cluster 1 **(Figure 3G**).

**Figure 3.**
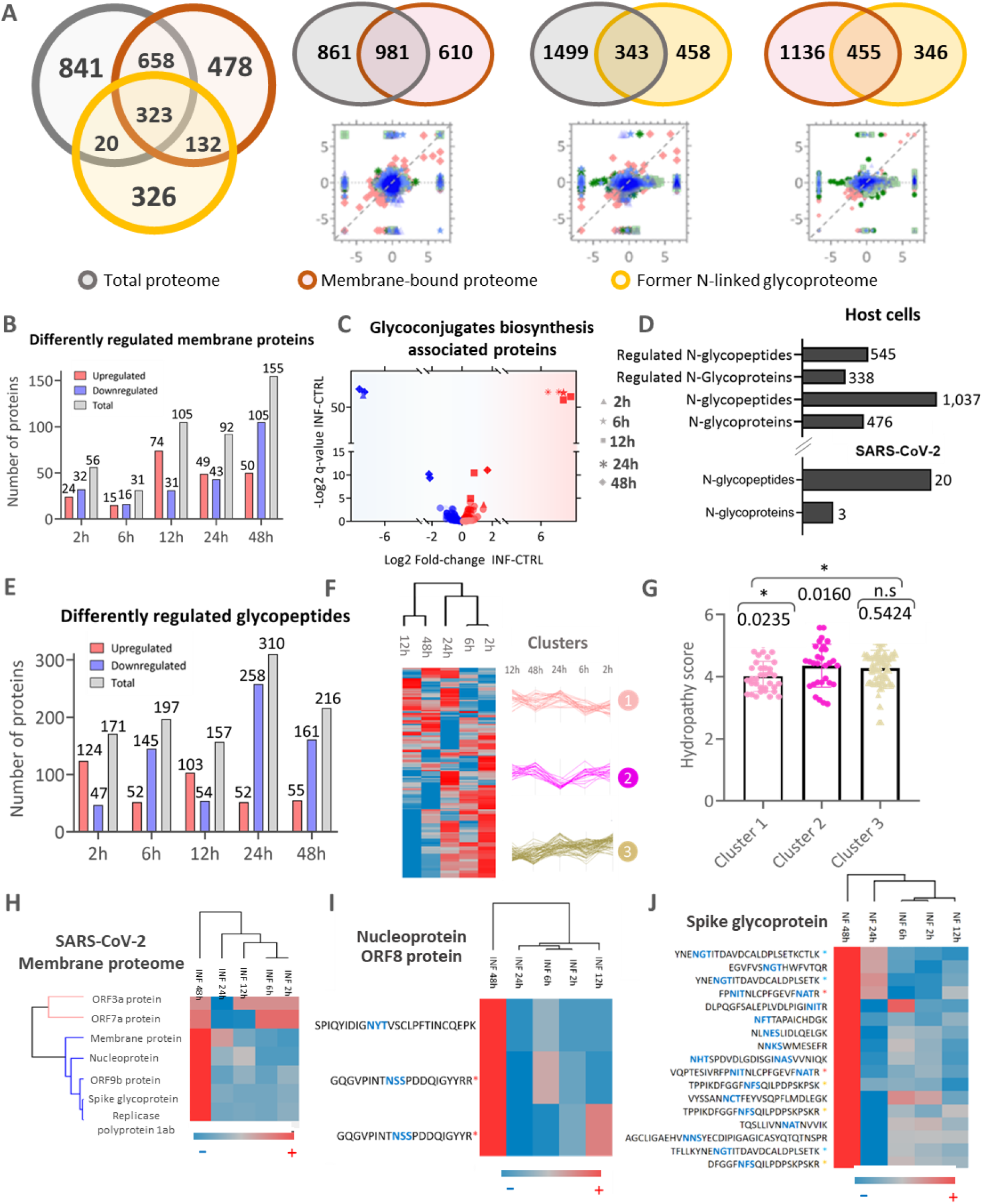
Membranome and former N-linked host cells upon SARS-CoV-2 infection. Venn diagram indicating common and exclusive proteins identified in the total proteome (gray), membranome (yellow), and N-deglycoproteome (orange) analysis. Scatter plots indicate the correlation between common proteins between the three datasets **(A)**; Vero cells proteins differentially regulated membrane proteins at 2, 6, 12, 24 and 48 hpi (q-value <0.05). Up, down and total regulated proteins are represented in red, blue and grey bars, respectively **(B)**. Regulation of membrane proteins associated with the glycoconjugates biosynthesis according to time of infection (**C**); Glycopeptides and glycoproteins identified in the host cells and SARS-CoV-2 **(D)**; Differently regulated N-deglycopeptides between the infected (INF) and control (CTRL) groups (q-value <0.05 **(E)**; Formerly N-linked glycopeptides differentially regulated in at least one comparison between infected (INF) and control (CTRL) groups **(F)**; Hydrophobicity score of clusters of differently regulated peptides in heatmap A (**G**); SARS-CoV-2 viral proteins identified together with host membrane (**H**); SARS-CoV-2 formerly N-linked glycopeptides mapped to nucleoprotein, replicase polyprotein 1a, ORF8 glycoprotein **(I)** and spike glycoprotein **(J)**. Blue sequences present N-glycosylation sequon.

Moreover, abundance of viral proteins identified together with host membrane increased sharply at 48 hpi, as expected due to viral replication (**Figure 3H**). We were also able to quantify 20 SARS-CoV-2 formerly N-linked glycopeptides, being 2 mapped to nucleoprotein, 1 to ORF8 protein and 17 to spike glycoprotein (**Figures 3I and J, Supplementary Data Set 2**). We modelled the spike protein and highlighted the identified glycosylation sites and their abundance change during infection (**Supplementary Figure 2**). Mapping the identified N-deglycopeptides associated to spike (P0DTC2) and ORF8 (P0DTC8) proteins surface illustrated possible sites crucial for their function (**Supplementary Figure 2**).

Functional enrichment analysis of host regulated membrane proteins showed cell death, stress response and transport related processes already modulated at 2 hpi. Processes and pathways related to post-translational modification and asparagine N-linked glycosylation were observed at 12 hpi, while apoptosis, protein folding and oxidative stress were among up-regulated processes at 24 hpi and 48 hpi (**Supplementary Data Set 3**).

By performing an integrated analysis of all MS-based approaches, we identified the formation of four clusters (**Figures 4A and B, Supplementary Data Set 3**), demonstrating again that ER-related processes and cell death are being regulated during infection (**Figure 4C and D, Supplementary Data Set 3**). ER-related processes were mostly regulated at intermediate time-points, similar to proteome findings. To visualize the regulated processes, we built protein networks for all clusters with regulated proteins found in the merged dataset. We found that Cell death, UPR and Response to endoplasmic reticulum stress shared common nodes (**Figure 4 E-I**).

**Figure 4.**
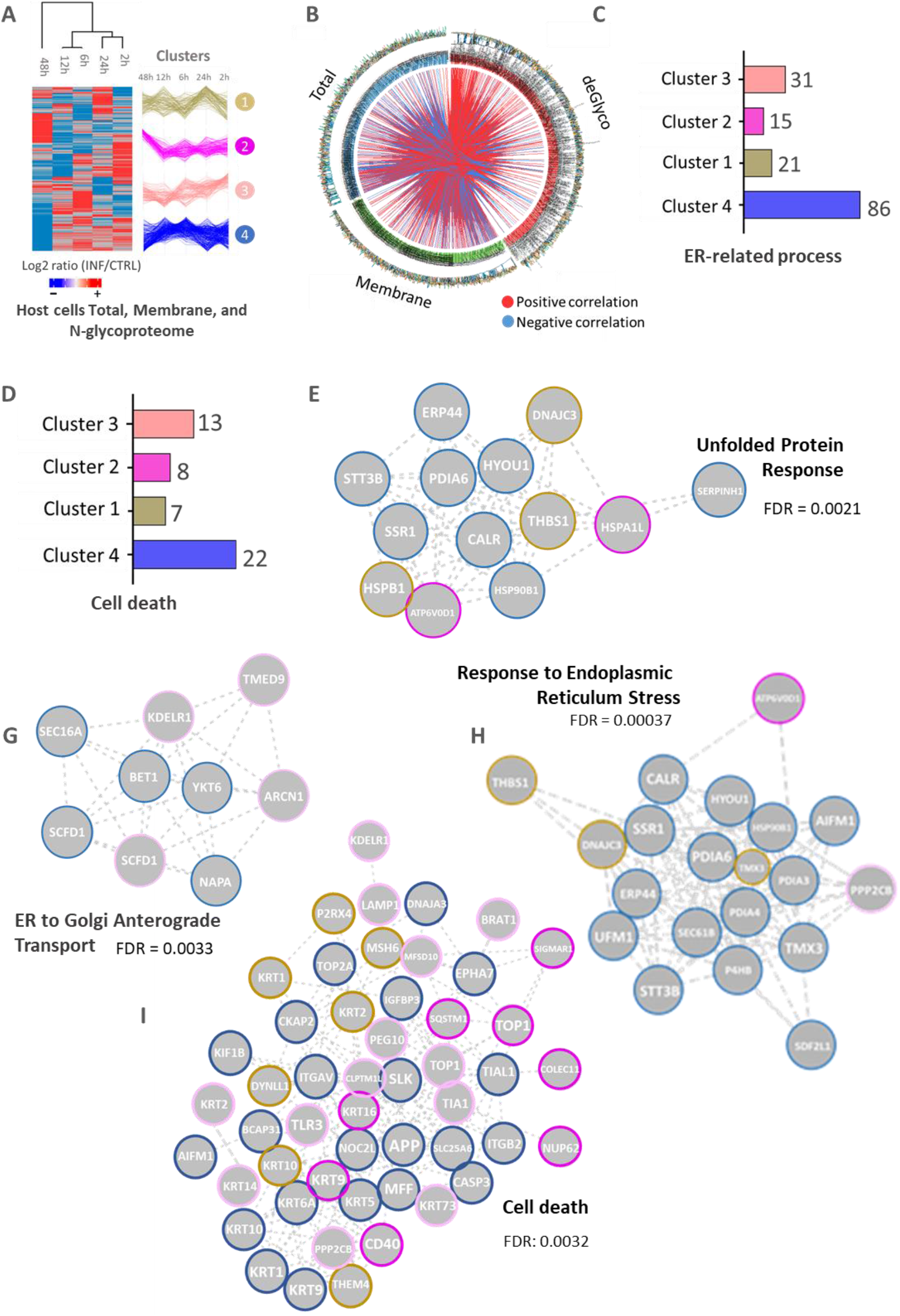
Integrative analysis of MS-based proteome, membranome and N-linked deglycoproteome. Differentially regulated host proteins in at least one group of a dataset **(A)**; Correlation map indicating the positive (red) or negative (blue) correlation between regulated proteins/peptides of different experimental approaches **(B)**; Proteins associated to Endoplasmic reticulum (ER) stress **(C)** and cell death **(D)** identified in the clusters of heatmap and respective enriched protein-protein interaction networks (q-value <0.05) **(E-I)**.

To further explore and confirm the effects of the viral infection, we performed immunoblotting analysis focusing on specific molecular pathways regulated in a time-course manner. In particular, we evaluated the activation of ER-stress, unfolded protein response (UPR), cell death and oxidative stress markers. This validation was performed in Vero CCL-81 monkey and Calu-3 human cell lines. Increased phosphorylation levels of PERK and eIF2α as well as protein levels of ATF4 were observed after 2h of viral infection presenting a peak at 6h in both cell lines tested (**Figures 5A-D, Supplementary Figures 3 and 4**). These results further confirmed that this UPR pathway was activated by the virus (**Figure 5A-D, Supplementary Figures 3 and 4**). In addition, higher levels of ATF6 and phosphorylated IRE1α were seen only after 48h of infection (**Figure 5E, Supplementary Figures 3 and 4**). Interestingly, phosphorylated IRE1α was not increased in Calu-3 cells (Supplementary Figure 3). The proteomic data have also detected higher levels of proteins related to apoptosis induction. Indeed, the western blot results demonstrated that CHOP, a protein linking UPR and apoptosis activation ^38^, presented increased levels upon 6h of viral infection only in the more susceptible Vero cells, indicating that apoptosis has been triggered in these cells by the virus (**Figure 5G, Supplementary Figures 3 and 4**). The detection of higher levels of cleaved caspase-3 clearly demonstrated apoptosis activation upon 48h of viral infection (**Figure 5H**).

**Figure 5.**
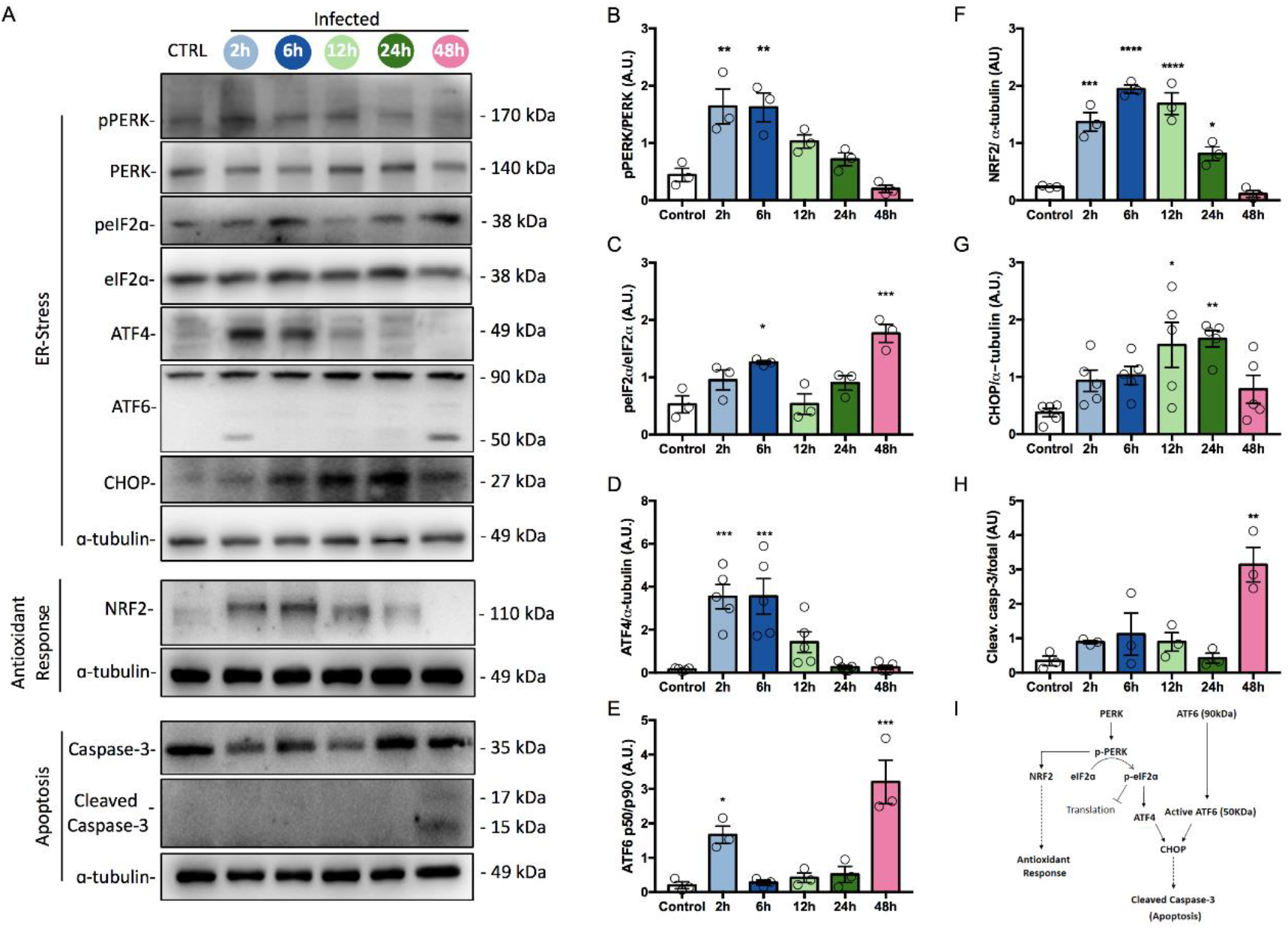
SARS-CoV-2 infection induces ER-stress, antioxidant response and apoptosis in Vero cells. Representative images of Western blots of ER-stress, antioxidant response and apoptosis proteins, as indicated **(A)**. The corresponding quantification of protein ratios of pPERK/PERK **(B)**, peIF2α/eIF2α **(C)**, ATF4/α-tubulin **(D)**, ATF6 p50/ ATF6 p90 **(E)**, NRF2/α-tubulin **(F)**, CHOP/α-tubulin **(G)** and Cleaved Caspase-3/Caspase-3 **(H)**. Schematic representation of ER-stress, antioxidant response and apoptosis pathways activated after SARS-CoV-2 infection **(I)**. Each dot represents an independent experiment (n?3 independent experiments; **** p<0.0001; *** p<0.001; ** p<0.005; *p<0.05 vs Control).

Additionally, we investigated PERK-NRF2 pathway axis to understand the increased antioxidant response observed in the infected cells. Higher levels of these proteins were observed after 6 and 12h of viral infection (**Figures 5F, 5H, Supplementary Figures 3 and 4**). Interestingly, the proteomic data evidenced significant decreasing levels of proteins displaying a function related to antioxidant response in Vero cells at the same time points shown in **Figure 5F** and **Supplementary Figures 3 and 4**. These data pointed at a generation of oxidative stress upon viral infection with the corresponding activation of early antioxidant response in the host cells. Viral infection has been shown to induce oxidative stress by ROS production to facilitate their replication in the host cell ^39,40^. In some cases, viruses have the ability to suppress the NRF2 pathway in their favor ^41^.

Since it is known that NRF2 can prevent cellular and tissue damage by decreasing the production of DAMPs (Damage-Associated Molecular Patterns) that are released by necrotic cells ^42^. In addition, oxidative stress generated by redox imbalance contribute to viral pathogenesis, resulting in a massive induction of cell death ^43^. Therefore, we decided to investigate some mechanisms of regulated cell death (RCD). Beside the caspase activation already described in Vero cells, we studied whether necroptosis and ferroptosis were also activated. For this purpose, the protein levels and/or the phosphorylation state of some components of these pathways were analyzed by western blot. Higher levels of MLKL phosphorylation were observed after 48h of viral infection in these cells indicating that part of cell death could be caused by necroptosis activation ^44^ (**Supplementary Figure 3**). This effect was not seen in Calu-3 cells. Higher levels of GPX4 at 6h of viral infection, in Vero and Calu-3 cells (**Supplementary Figure 3 and 4**), could indicate that ferroptosis was not being activated, because GPX4 may be part of the antioxidant mechanism activated by the PERK-NRF2 pathway, since GPX4 is also an established NRF2 transcriptional target ^45^. Overall, these results indicate that at least two cell death regulated pathways are being activated by viral infection in Vero cells. Unlike what was observed in Vero cells, Calu-3 cells presented no changes in cleaved caspases or MLKL phosphorylation levels upon viral activation until the last time point studied. These results led us to conclude that different cells may display different kinetics in cell death signaling pathways activation. This could be related to the existence of stronger homeostatic responses being triggered to avoid cell death. Indeed, previous results from our group have shown that Calu-3 cells exposed to viral infection start show signs of cell death only after 72h (data not shown).

To access further translational aspects of our *in vitro* findings, we re-analyzed transcriptome data obtained from lung autopsies of eight patients who died as a result of COVID-19 (**Figure 6A**) and the respective controls ^37^. Using a dedicated pipeline to reprocess the data with higher stringency in the statistical test, we identified 1,398 regulated transcripts, being 636 up-regulated and 762 down-regulated (**Figure 6B**). PCA analysis showed diverse transcriptome profile between COVID-19 patients and controls. (**Figure 6C**). We observed that differentially regulated transcripts were involved in several processes linked to ER stress, such as cell death, chaperone-mediated folding, ‘de novo’ protein folding, protein localization to ER, programmed cell death, and protein folding, confirming the proteomic data (**Figures 6G and H**). Mapping ER-stress transcripts and proteins in clinical specimens from patients infected with SARS-CoV-2, it was possible to identify that RCN3, UCHL1, and ERO1A are upregulated in the lung at the level of transcript and proteome ^37^. Moreover, we found 51 up-regulated and 45 down-regulated confirming the alteration of the host glycosylation biosynthetic machinery upon SARS-CoV-2 infection (**Figure 6F)**. Interestingly, hierarchal clustering analysis showed that the infected samples 1, 3, and 4 had a distinguished pattern of up-regulated ER-related transcripts (**Figure 6D**). It is worth to mention that the average survival time after being hospitalized was significantly higher in these three patients compared to the others (**Figure 6E**). These data confirm the regulation of ER-stress proteins during SARS-CoV-2 infection.

**Figure 6.**
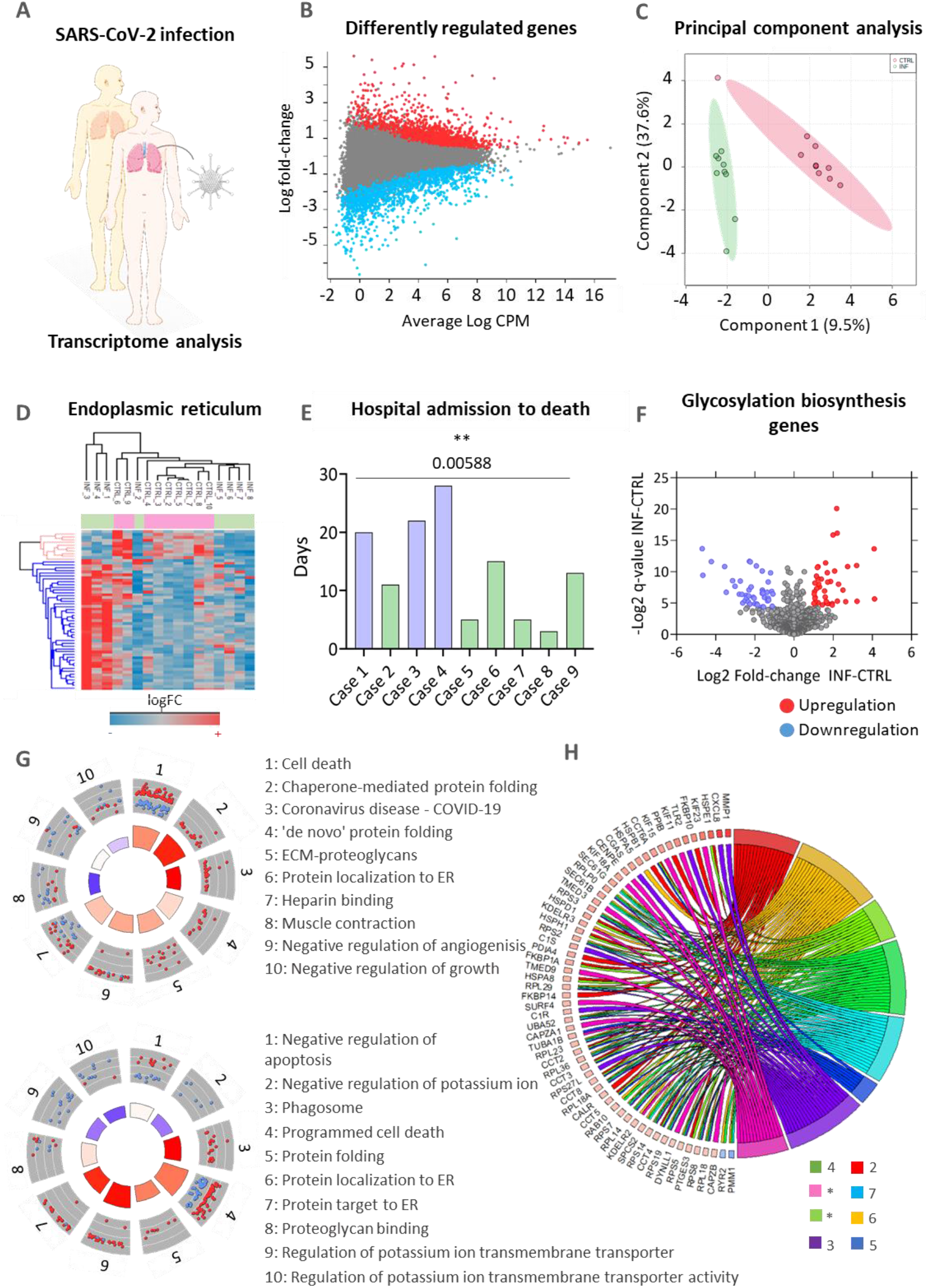
RNA-seq reanalysis data including samples of the lung of patients infected with SARS-CoV-2 and healthy samples from cancer donors (A), indicating the Differently regulated genes (B); Principal component analysis (C); Heatmap for differently regulated genes that are located in the endoplasmic reticulum (ER) (D); Hospital admission to death in days (E); Genes associated with the glycosylation biosynthesis alteration (F) and Gene ontology analysis (G-J). The (*) corresponds to Golgi-to-ER retrograde transport and the Establishment of protein localization to endoplasmic reticulum pathways.

Taken together, our data indicated that SARS-CoV-2 infection modulates glycoconjugates biosynthetic machinery changing the host global protein glycosylation profile. Additionally, ER stress induced in infected cells activates PERK-eIF2α-ATF4-CHOP UPR pathway finally leading to apoptosis induction. Cell death might also occur by necroptosis, linked to antioxidant response and activation of PERK-NRF2 pathway and MLKL phosphorylation. These *in vitro* phenomena were also observed in human lung biopsies of COVID-19 patients, indicating a role of ER protein modulation and survival time.

## DISCUSSION

SARS-CoV-2 hijacks several host machineries to control immune reaction ^46^, viral protein translation ^47^, viral genome packing into nascent viral particles as well as support the release of mature virus particles. Host cellular machineries are redirected to synthesize and remodel viral proteins through post-translational modifications such as proteolytic cleavages, disulfide bridges formation, phosphorylation, ubiquitination and glycosylation ^48–53^. Thus, in this study we sought out which processes and pathways could be changed by the viral infection. Our proteomics approach yielded a total of 2778 proteins being 1842, 1591 and 801 proteins identified in the total, membrane and glycosylated proteome, respectively. Even if the number of total identified proteins is comparable to other studies using similar technological platforms, we analyzed different time-points, cell lines and MOI. Grenga et al. (2020) identified 3220 host proteins and 6 SARS-CoV-2 proteins over 5 time-points evaluated (1, 2, 3, 4 and 7 days) and two MOI (0.1 and 0.001) ^54^. Stukalov et al. (2021) identified a total of 5862 proteins in ACE2-expressing A549 cells infected with SARS-CoV-2 and SARS-CoV over three time points (6, 12, 24 hpi); concerning only regulated proteins, they found a total of 272 ^48^, while we identified 443 regulated proteins using only Vero cells infected with SARS-CoV-2 over five time points. It should be noted that we used an earlier (2 h) and later (48 h) time point that influence the number of regulated proteins. Bojkova et al. (2020) identified over 7,000 proteins using Caco-2 infected cells at four time-points (2, 6, 10 and 24 hours) being over 3,400 regulated proteins ^55^. Using two different MOI (0.1 and 3), Zecha et al. (2020) identified 7,287 proteins and approximately 1,500 regulated host proteins in SARS-CoV-2 infected Vero cells in a single time-point (24 hpi) ^56^.

Our results pointed at host proteome remodeling upon viral infection, consisting of protein global downregulation in all evaluated time points, except at 6 and 48 hpi. Such pattern was also recently reported in the total proteome analysis performed by Stukalov and collaborators (2021). Although they did not analyze the proteomic profile at 48 hpi, they observed that proteins were mostly down-regulated at 12 and 24 hpi but being up-regulated at 6 hpi ^48^. Since it has already been shown that viral replication starts at 6 hpi ^57,58^, the observed global protein up-regulation at 6 hpi could imply an initial response from the cell, followed by protein inhibition due to viral influence until 48 hpi, when cell death events are most prevalent. This remodeling was supported by PCA analysis, which showed a clear time-dependent separation.

Among the identified proteins, we found 8 viral proteins in all time-points. Compared with the work in Caco-2 cells performed by Bojkova et al. (2020) ^58^, we did not identify the non-structural protein 8, and instead of identifying the replicase polyprotein a, we have identified the replicase polyprotein 1ab. Our data have confirmed part of the data published by Davidson et al. ^59^using Vero cells. Indeed, we have identified the non-structural protein 9b but not proteins 8 or 9a. The viral proteins N, S, M, ORF1ab, ORF3a and ORF7a identified in this study were also found in the recent work of Grenga and collaborators (2020) ^54^.

Regarding regulated biological processes and pathways, our results showed similarities when compared to other proteomics studies. A cluster analysis of SARS-CoV-2 infected Vero cells over a period of seven days indicated that membrane trafficking, protein pre-processing in the ER, clathrin-mediated endocytosis, vesicle-mediated transport, and viral life cycle were enriched during the infection ^54^. Similarly, we have reported membrane trafficking, pathways related to the viral life cycle, and post-translational protein modification enriched in the regulated protein dataset. As already shown in infected Caco-2 cells, our pathway analysis has also identified that upon viral infection, TCA and respiratory electron transport as well as the carbohydrates metabolism processes were modulated. Other common regulated processes observed in the literature are autophagy, IFN-α/β induction or signaling, cell adhesion, and extracellular matrix organization ^48,56^, being the regulation of IFN-α/β pathway extensively explored as a drug target for viral replication inhibition ^48^. Besides similar processes and pathways during SARS-CoV-2 infection, we have focused on the effects of viral replication in the ER-stress and UPR in a time-dependent manner. Through hierarchical clusters, we identified 3 main clusters that showed the effects of the viral infection in a time-dependent manner. Among the biological processes identified in the clusters, asparagine N-linked protein glycosylation, response to stress, unfolded protein response and post-translational protein modification were increased in the intermediate time-points, but decreased at 48 hpi.

Glycosylation of viral proteins is an important process that regulates viral assembly and infectivity. The structural and functional role of glycosylation in the SARS-CoV-2 spike protein has been widely investigated ^52,60–66^. Of note is the fact that 35% of the SARS-CoV-2 spike glycoprotein contains carbohydrate moieties, which have profound influence on the viral infectivity, susceptibility to antibody neutralization ^67,68^. The N-linked glycosylation sites of spike proteins have been related to alterations in its open or closed state thus interfering in its capacity to bind to the receptor ^69^. We found 17 formerly N-glycopeptides and 14 glycosylation sites in the Spike glycoprotein. There are 22 potential glycosylation sites in the SARS-CoV-2 Spike protein and the number of reported occupied sites range between 17 and 22 ^51,63,70,71^. The processing of the spike glycoprotein through the ER and Golgi compartments represents an important step in controlling the virion assembly and inhibition of the N- and O-glycan maturation, which has been shown to interfere with virulence ^70,72–75^. Additionally, we mapped the N-linked glycosylation site of ORF8 protein of SARS-CoV-2. This accessory protein has less than 20% identity with the same protein in SARS-CoV, highlighting divergencies between the two viruses ^76^. Although this protein does not appear to be essential for viral replication, it has been shown to disrupt IFN-I and promote MHC-I downregulation ^77,78^. ORF8 contains a signal peptide for ER import and interacts with several proteins within the ER. In this study, we have mapped one N-linked glycosylation site at N78. This site is close to a SARS-CoV-2-specific sequence YIDI^76^, that has been reported to be involved in noncovalent dimerization ^36^. Antibodies against ORF8 were identified as serological markers of acute, convalescent and long-term response to SARS-CoV-2 infection ^79^. Therefore, it would be relevant to evaluate the role of site-specific ORF8 glycosylation in antibody neutralization.

The fact that asparagine N-linked glycosylation was enriched in the biological processes regulated during viral infection may indicate that the continuous translation of viral glycoproteins is overwhelming the glycosylation machinery capacity, increasing the number of proteins with aberrant glycosylation. This dysregulated process could contribute with ER stress, since it can increase protein misfolding ^80,81^. Proteins related to asparagine N-linked glycosylation were reported among the top 10% of proteins following viral gene expression ^58^. Thus, indicating a metabolic challenge for the host glycosylation machinery promoted by viral infection. A recent study has demonstrated that N-glycosylation inhibitors were able to reduce SARS-CoV-2 infection in Vero and HEK293^ACE-2^ cells. Moreover, genetic ablation of this pathway using siRNAs and virions presenting N-glycosylation defects also reduced the infection rate ^82^. Since we found that five enzymes involved in the glycosylation biosynthesis (CHST12, CHST14, B4GALT3, GCNT1 and MGAT2) were mainly up-regulated in the intermediate time-points, but down-regulated at 48 hpi, our data showed a complete remodeling of the N-linked protein glycosylation process. Recently, the downregulation of this process was also evaluated by the targeting of 88 host glycogenes by siRNAs in a study on the secretion of hepatitis B surface antigen (HBsAg) and HBV DNA, which showed that targeting CHST12 reduced the HBV DNA levels by >40% in EPG2.2.15.7 cells ^83^. This further support the hypothesis that viral the viral infection can modulate the host glycosylation machinery.

Aberrant glycosylation can interfere in protein folding (PMID: 24609034). The effects of viral infection on protein folding were also observed in the functional enrichment analysis of the differentially abundant proteins identified in this study. Indeed, at 6 hpi we observed an increase in ER stress caused by misfolded or unfolded proteins with the up-regulation of UPR. Moreover, we confirmed the activation of ER stress and UPR in SARS-CoV-2-infected Vero and Calu-3 cells using western blotting. Several evidences suggest that ER stress and UPR activation are the main contributors to the pathogenesis of various diseases including viral infections ^84^. Recent SARS-CoV-2 host interactome has been performed in HEK293, human bronchial epithelial 16HBEo- and A549 cells ^6,48,85^. Proteins related to ER stress such as thrombospondin-1, GRP78, DJB11, calnexin and F-box only protein 2 were found to interact with the spike protein ^86,87^. In addition, other SARS-Cov-2 proteins were found to interact with proteins involved in ER protein quality control, ER morphology and protein glycosylation ^6^. Cell surface GRP78 was identified to interact with the Middle East respiratory syndrome coronavirus spike glycoprotein and increase the viral entry ^88^. Furthermore, SARS-CoV S glycoprotein was found to bind calnexin and increase its infectivity by modulating the maturation of the glycans ^89^. Another host-virus protein-protein interaction analysis also pointed ER stress as being one of the pathways most affected by the SARS-CoV-2 proteins ^90^. SARS-CoV 3a protein has been shown to be able to induce ER stress by activating PERK pathway resulting in increasing levels of eIF2α phosphorylation and ATF4 protein level, which finally promoted the synthesis of CHOP and increased Huh7 cells apoptosis ^91^. It is important to note that the authors have not observed signs of ATF6 signaling pathway activation ^91^. Another study with SARS-CoV showed that the suppression of the spike protein inhibits the up-regulation of BiP and GRP94 chaperones, which are targets of PERK-eIF2α-ATF4 pathway activation. Moreover, the authors reported the inhibition of this pathway promoted a decrease of these chaperones, pointing once again that UPR modulation by the virus could facilitate the infection process ^92^. The stress-responsive heat shock protein gene HSP90AA1 was reported to be induced in H1299 and Calu-3 cells during infection, adding more evidence of ER stress occurring upon SARS-CoV-2 replication ^93^. The activation of CHOP due to stress can induce the expression of BIM, linking ER stress induction with apoptosis activation ^94^. Moreover, ATF4 and CHOP can also activate the translation of genes related to translational components which will enhance protein synthesis in the cell, causing an increase in ROS production and consequently cell death ^95^.

It has been shown prolonged ER stress can activate apoptosis pathway, which will conclude with the assembly of the apoptosome and caspase-3 activation ^96^. We observed that apoptotic-related processes were mainly modulated in the late time events, indicating that cell death may be more frequent at 48 hpi. Additionally, the enriched pathway analysis showed that these processes started at 12 hpi and remained active at 24 hpi. Although ER stress is predominantly the main cause of stress observed in this study, viruses have been shown to induce oxidative stress by ROS production in to facilitate their replication in the host cell ^39,40^. Viral infections can induce the release of pro-oxidant cytokines such as the tumor necrosis factor (TNF), which lately will produce the hydroxyl radical OH ^97^. A study in Huh-7 cells indicated that Ca^2+^ released from the ER as a consequence of the human hepatitis C virus-induced ER stress will lead to an increase Ca^2+^ upload by the mitochondria, where it will promote the generation of higher ROS levels and the consequent increase in oxidative stress ^98–101^.

Besides caspase activation, we found increased phosphorylation of MLKL at 48 hpi indicating a possible contribution of necroptosis induction upon viral infection. This dual mode of cell death mechanism has been reported in infected HFH4-hACE2 transgenic mouse model, Calu-3 cells and in postmortem lung sections of fatal COVID-19 patients ^102^.

Since our study clearly points towards protein folding, ER stress and UPR modulation, which lead to cell death in later events, we sought if the effects of viral infection in these processes could also be seen not only *in vitro* models but also in human tissue biopsies. Our re-analysis of the post-mortem lung transcriptome of COVID-19 patients showed that processes related to protein folding and cell death are being regulated, which are in accordance with proteomics analysis showing increased levels of ER stress and UPR modulation. The processes identified in our re-analysis were not explored by the authors of the original work as their results focused on neutrophil activation and neutrophil-mediated immunity, extracellular traps and extracellular structure organization ^37^. Interestingly, ER stress related pathways and processes are not well explored in transcriptomic studies, being processes and pathways related to immune response and inflammation more commonly found in the literature, such as modulation of cytokine-mediated signaling pathway, interferon signaling, TNF-signaling, and interleukin-mediated signaling ^54,103–105^. Nevertheless, the enrichment analysis of these works often contains processes and pathways related to proteins transport and localization to ER. For example, the transcriptional response of hACE2 receptor-transduced A549 and Calu-3 cell lines to SARS-CoV-2, MERS-CoV, or influenza A virus (IAV) infections focused on the autophagy pathway and mitochondrial processes, but UPR modulation was observed in the A549 lung epithelial cell line ^106^. Interestingly, the authors compared the A549 cells infected with SARS-CoV-2 and IAV and found UPR modulated only in SARS-CoV-2 infection ^106^. A recent study has shown that recombinant expression of SARS-CoV-2 spike protein in HEK293T cells induces ER-stress and UPR activation ^107^. The increased expression of GRP78 and phosphorylated eIF2α was reported together with increased LC3 II at 24 hours post spike transfection ^107^. Treatment of transfected cells with UPR modulators reduced the ER stress levels ^107^. However, this study did not provide a kinetic and comprehensive measurement of ER-stress and UPR activation, as reported here.

Our results corroborate recent findings on the identification of serum ER stress markers (GRP78 and phosphorylated PERK) in the lungs of COVID-19 patients with severe complications ^108^. Proteomic analysis of multiple organs from patients infected with SARS-CoV-2 showed that, in addition to the lungs, there is an increase in ER stress in the renal cortex and liver cells. RCN3 is a protein located in the ER lumen that acts in the remodeling during lung injury ^109^. RCN3-deficient mice has been shown to result in increased ER stress and apoptosis ^109,110^. The ERO1A protein is a CHOP-activated oxidase that promotes ER hyperoxidization and affects the activation of CHOP-dependent apoptosis by stimulating the IP3R1 1,4,5-triphosphate inositol receptor ^111,112^. The UCHL1 protein performs important functions related to protein degradation ^113^. In addition, it was observed that UCHL1 levels influence cell homeostasis under normal conditions of growth and oxidative stress ^114^. The transcriptome of silenced cells for UCHL1 showed downregulation of genes associated with proteasome activity and upregulation of genes linked to ER-stress ^115^. Moreover, we observed that samples from COVID-19 patients with higher expression of ER transcripts were associated to longer survival period.

Taken together, this study provides a time-resolved and large-scale characterization of the total, membrane and glycoproteome of SARS-CoV-2-infected Vero CCL-81 cells. The modulation of specific processes including viral and host protein glycosylation, ER-stress and UPR were validated using western blotting and reanalysis of transcriptomic data of human clinical specimens. These data highlight the importance of ER-stress and UPR modulation as a host regulatory mechanism during viral infection and could point to novel therapeutic targets.

## Acknowledgments

The work was supported by grants and fellowships from FAPESP (2018/18257-1, 2018/15549-1, 2020/04923-0 to GP; 2016/04676-7 to AFS; 2019/09517-2 to GSA; 2017/24769-2 to RRGM; 2016/20045-7 and 2020/06409-1 to ELD), from Coordenação de Aperfeiçoamento de Pessoal de Nível Superior – Brasil (CAPES) (Código de Financiamento 001 to LR-F; 88887.131387/2016-00 to DBA) and from Conselho Nacional de Desenvolvimento Científico Tecnológico (CNPq) (“bolsa de produtividade” to GP).

Professor Suely Kazue Nagahashi Marie from FM-USP is acknowledged for advice during manuscript writing and all the support.

## Author Contributions

GP conceived the idea. LR-F and GP designed the experiments. LR-F, LCL, CBA and GP prepared the samples for proteomics analysis. LR-F, LCL, JMdS and GP analyzed the mass spectrometry data, performed bioinformatic analyses and wrote the manuscript. RRGM, DBA, DBLO and ELD performed the viral infection and RT-qPCR. CW and CRFM assisted on data interpretation. VdMG, AFS, GSA and LL performed and analyzed the western blotting. All authors contributed in editing the manuscript and approved the final version.

