## Supplementary data for "SARS-CoV-2 activates ER stress and Unfolded protein response"

**SUPPLEMENTARY MATERIAL**

**Supplementary Data Sets**

**Supplementary Data Set 1**. Data obtained from the analysis of the total proteome, indicating the Proteome discoverer search result, differently regulated proteins at 2h, 6h, 12h, 24h, 48h; clusters; enriched results for Vero cells clusters; principal component analysis (PCA); subcellular location analysis; mapped glyco-genes; subcellular location of mapped glyco-genes; and gene ontology result for upregulated/downregulated proteins at 2h, 6h, 2h, 24h, and 48h.

**Supplementary Data Set 2**. Data obtained from the analysis of the membrane and deglycoproteome, indicating the Proteome discoverer result; identified N-glycopeptides; differently regulated proteins (membrane) at 2h, 6h, 12h, 24h, and 48h; Differently regulated glycopeptides at 2h, 6h, 12h, 24h, and 48h.

**Supplementary Data Set 3**. Data obtained from the analysis of the integrated total, membrane, and deglycoproteome indicating the clusters (datasets) and clusters gene ontology.

**Supplementary Data Set 4**. RNAseq data re-analysis, indicating patient information; pre-processed matrix (filtered and normalized); differently regulated genes (|FC> 1| and q- value <0.05); mapped genes associated with glycosylation biosynthesis; results of the enrichment analysis; ER proteins.

**Supplementary Figures**

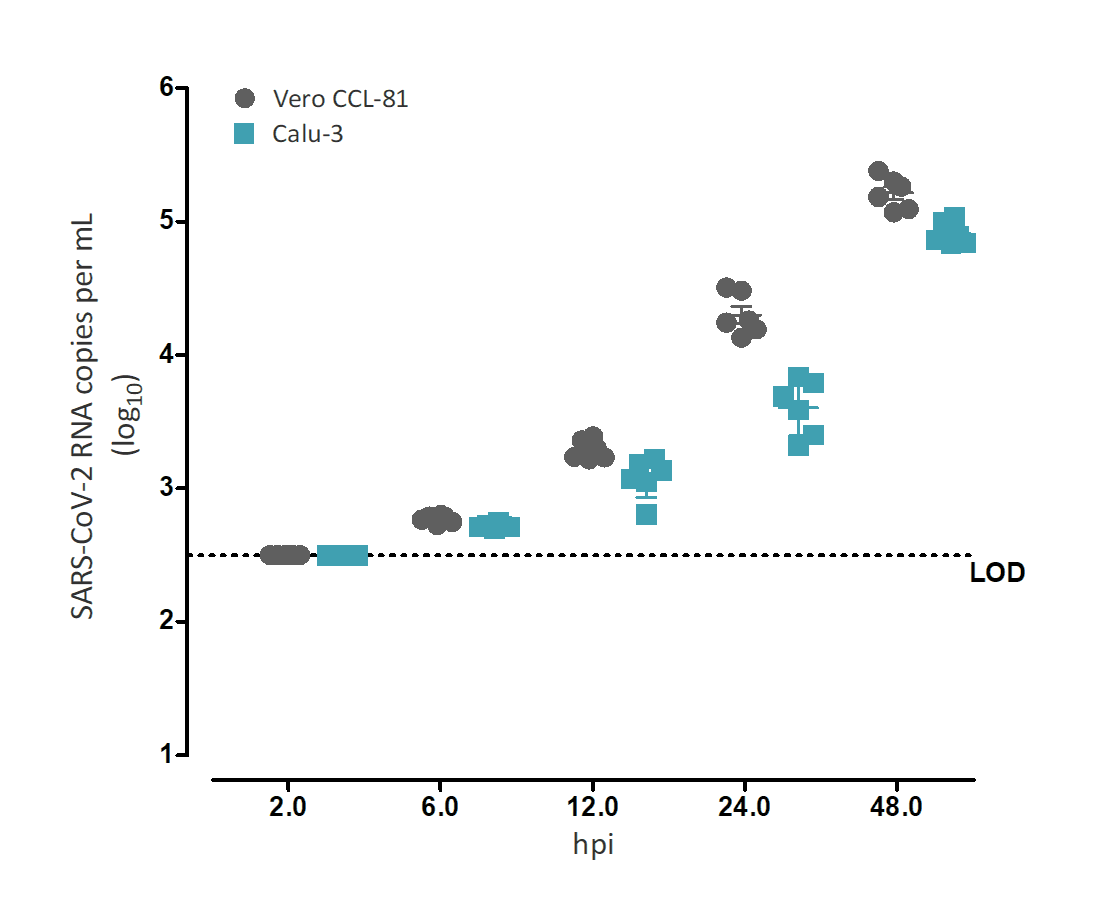

**Supplementary Figure 1** Quantification of viral infection progression over time

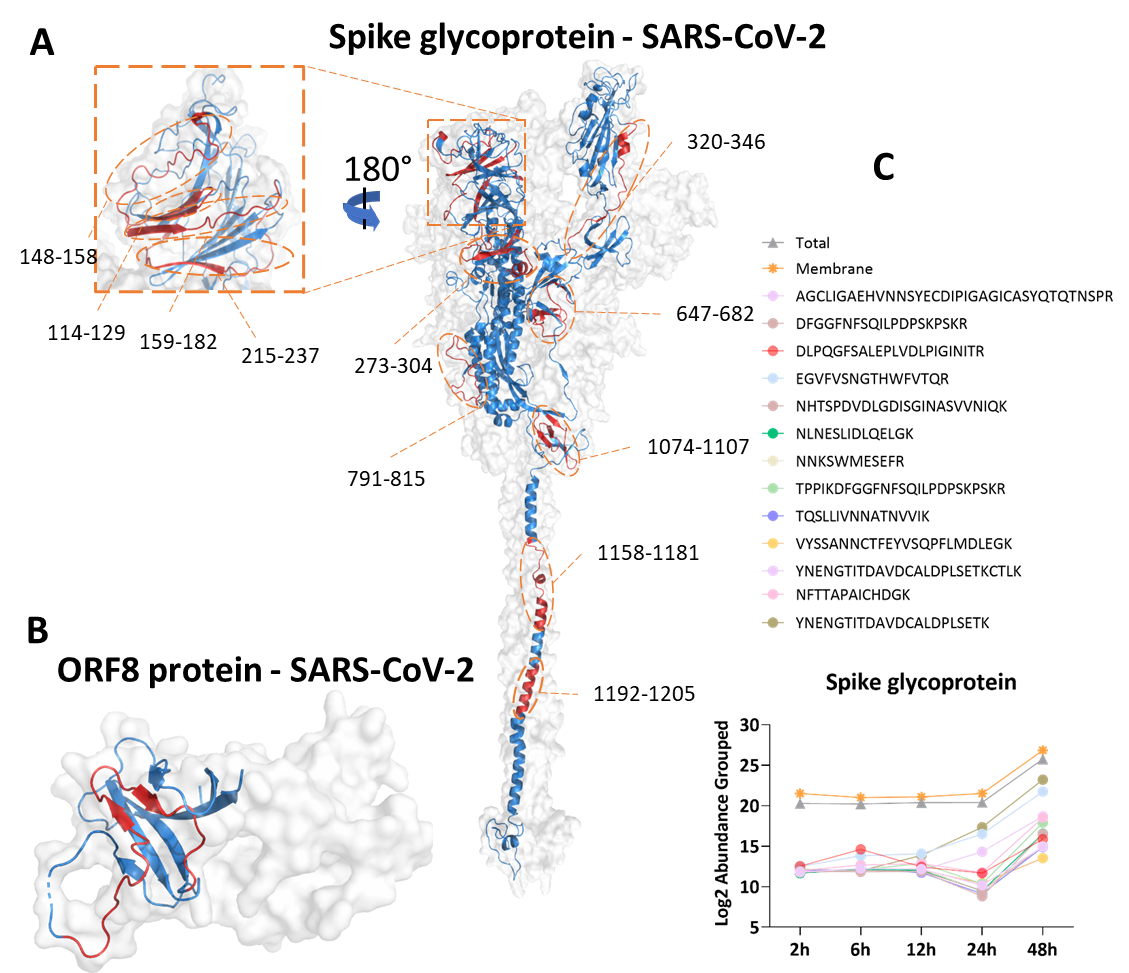

**Supplementary Figure 2**: Structural representation of P0DTC2 (A) and P0DTC8 (B) surface and backbone structure with the identified peptides highlighted in red. Chain A backbone is represented in light-blue while other monomers where omitted. Overlapping identified peptides regions where combined for simplicity; Quantitative proteome of P0DTC2 in total, membrane, and deglyco datasets (C).

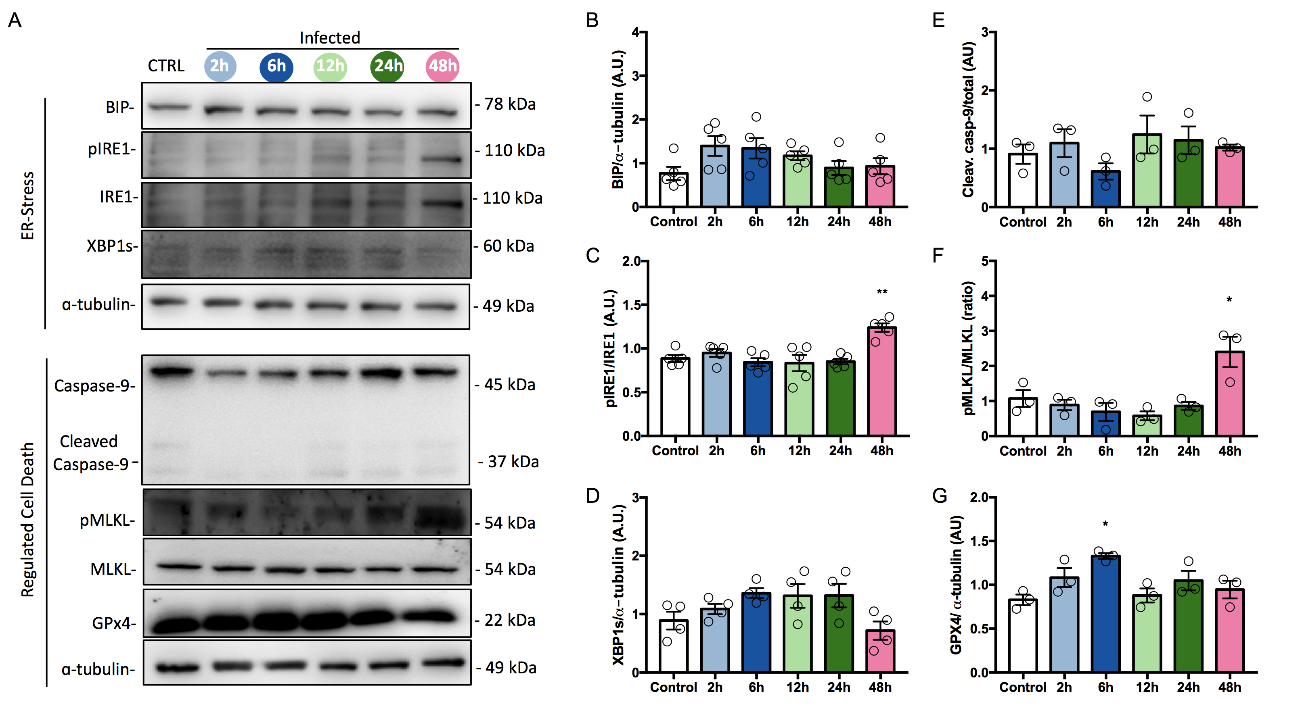

**Supplementary Figure 3**: SARS-CoV-2 infection induces ER-stress and necroptosis in Vero cells. (A) Representative images of Western blots of other branches of ER-stress pathway and regulated cell death, as indicated. The corresponding quantification of protein ratios of (B) BIP/ɑ-tubulin, (C) pIRE1/IRE1, (D) XBP1s/ɑ-tubulin, (E) Cleaved Caspase-9/Caspase-9, (F) pMLKL/MLKL and (G) GPX4/ɑ-tubulin. Each dot represents an independent experiment (n≥3 independent experiments). **** p<0.0001; *** p<0.001; ** p<0.005; *p<0.05 vs Control.

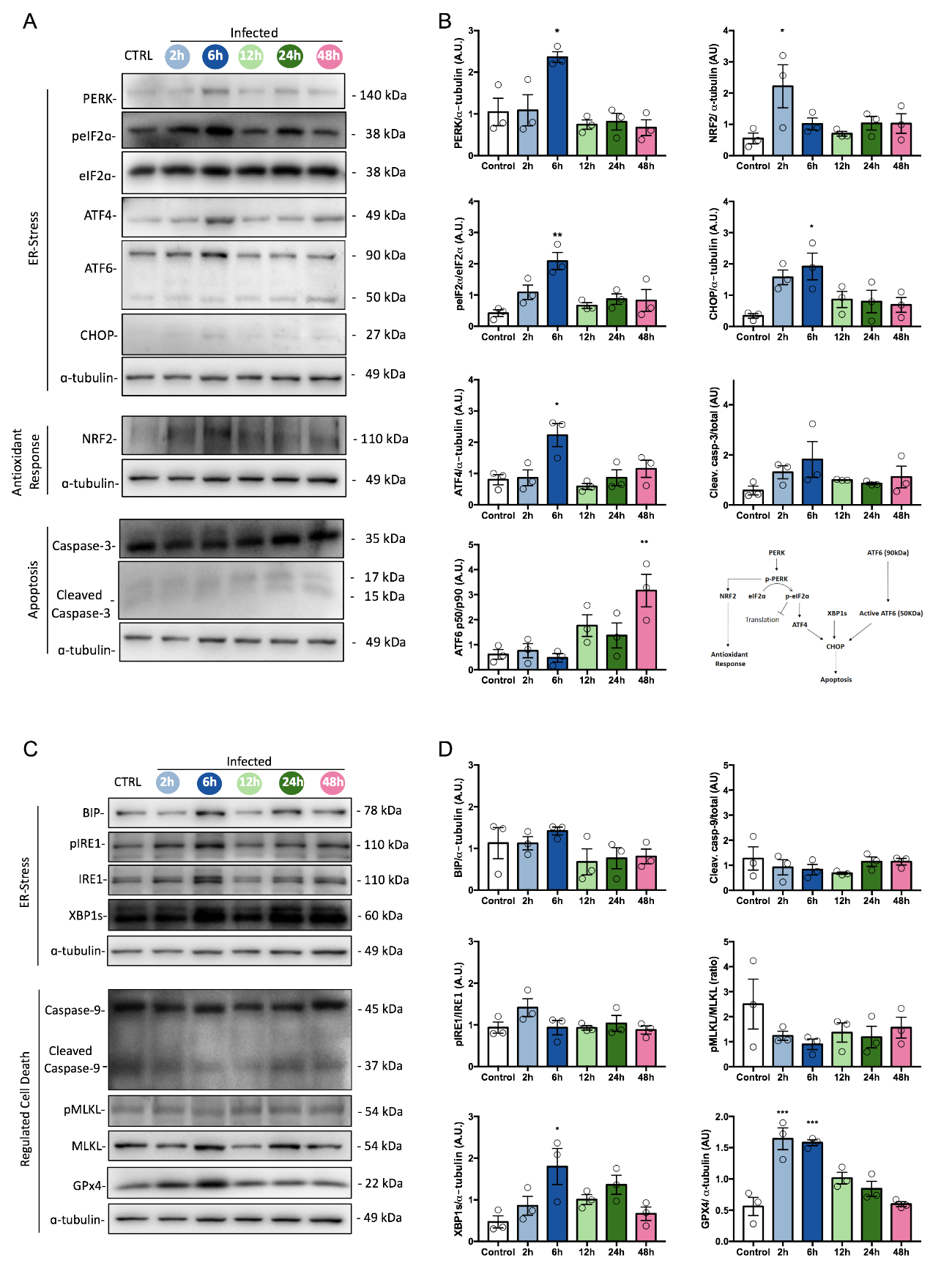

**Supplementary Figure 4**: SARS-CoV-2 infection induces ER-stress and antioxidant response in Calu-3 cells. (A) Representative images of Western blots of ER-stress, antioxidant response and apoptosis proteins, as indicated. (B) The corresponding quantification of protein ratios of pPERK/PERK, peIF2ɑ/eIF2ɑ, ATF4/ɑ-tubulin, ATF6 p50/ ATF6 p90, NRF2/ɑ-tubulin, CHOP/ɑ-tubulin and Cleaved Caspase-3/Caspase-3 and schematic representation of ER-stress, antioxidant response and apoptosis pathways activated after SARS-CoV-2 infection. (C) Representative images of Western blots of other branches of ER-stress pathway and regulated cell death, as indicated. (D) The corresponding quantification of protein ratios of BIP/ɑ-tubulin, pIRE1/IRE1, XBP1s/ɑ-tubulin, Cleaved Caspase-9/Caspase-9, pMLKL/MLKL and GPX4/ɑ-tubulin. Each dot represents an independent experiment (n≥3 independent experiments). **** p<0.0001; *** p<0.001; ** p<0.005; *p<0.05 vs Control. (I) Schematic representation of ER-stress, antioxidant response and apoptosis pathways activated after SARS-CoV-2 infection.
